# Rapid Shift Toward Pulsed Field Ablation and Precision Risk Stratification in High-Impact Atrial Fibrillation Research

**DOI:** 10.64898/2026.09.11.751096

**Authors:** Zheng Su, Tinsley Li

## Abstract

Conventional bibliometrics rely on lifetime citations, obscuring immediate shifts in cardiovascular research paradigms. To track emerging trends in atrial fibrillation management, we performed a comparative bibliometric analysis of the 50 highest-cited original research articles per year in OpenAlex topic T10065 across consecutive 2023 (Class of 2025) and 2024 (Class of 2026) publication cohorts. Articles were ranked using a fixed 18-month post-publication citation window, and extracted concepts were normalized into 719 canonical topics and 32 parent themes using large language model curation. Concept frequency tracking demonstrated a swift technological shift, with pulsed field ablation showing the largest topic frequency increase (+0.08, from 0.30 to 0.38) to become the leading canonical topic in 2024, displacing conventional thermal pulmonary vein isolation (-0.18, 0.40 to 0.22). Simultaneously, stroke prevention focus shifted toward refined predictive modeling, with increases in Risk Stratification and Predictive Models (+0.08, 0.30 to 0.38) and CHA2DS2-VASc scoring (+0.08, from 0.10 to 0.18). High-impact atrial fibrillation research is rapidly pivoting toward non-thermal ablation safety profiling and precision risk stratification, highlighting the utility of fixed-window concept mining for capturing real-time scientific evolution. Online explorer of the result is available at https://pri.pepkio.com

## INTRODUCTION

Atrial fibrillation represents the most common sustained cardiac arrhythmia encountered in clinical practice, posing a formidable global public health burden that currently affects over 50 million individuals worldwide^1^. Driven by population aging, increasing cardiovascular risk factor prevalence, and enhanced diagnostic monitoring, the incidence and prevalence of atrial fibrillation continue to rise globally, contributing to elevated risks of ischemic stroke, heart failure, cognitive impairment, and cardiovascular mortality^2^. To mitigate this growing disease burden, clinical management strategies have historically pivoted around two central clinical priorities: thromboembolic risk reduction through oral anticoagulation therapy and rhythm control via invasive catheter ablation^3,4^ Over the past two decades, extensive clinical trial evidence and refinement of interventional techniques have firmly established catheter-based pulmonary vein isolation and guideline-directed stroke risk stratification as the twin pillars of contemporary atrial fibrillation care^5,6.^

In interventional electrophysiology, pulmonary vein isolation has long relied on thermal ablation modalities, predominantly radiofrequency energy and cryoballoon ablation, to achieve durable pulmonary vein isolation^7,8^. However, the therapeutic landscape is undergoing an unprecedented shift with the clinical introduction of non-thermal pulsed field ablation. By utilizing high-voltage electrical fields to induce irreversible tissue-selective electroporation, pulsed field ablation offers rapid lesion formation while minimizing thermal injury to adjacent non-cardiac structures such as the esophagus and phrenic nerve ^9-11^.

In parallel, stroke prevention paradigms are rapidly expanding beyond traditional binary antithrombotic management. Recent research has re-evaluated established clinical risk scores, comparing the classic CHA2DS2-VASc score against non-sex CHA2DS2-VA models to optimize antithrombotic decision-making12. Furthermore, clinical attention is extending toward expanding anticoagulation indications in subclinical atrial fibrillation^13,14^ and developing novel factor XI and XIa inhibitors aimed at decoupling stroke prevention efficacy from major bleeding risk^15,16^.

Despite the rapid influx of clinical trials, real-world registries, and technological innovations across these domains, capturing how scientific attention redistributes across emerging paradigms in real time remains a critical challenge. Conventional bibliometric methodologies and literature reviews predominantly rely on cumulative lifetime citation counts. However, lifetime citation metrics inherently favor older, established publications and introduce a substantial time-lag bias that obscures immediate post-publication shifts in scientific interest17.

Furthermore, traditional literature mapping relies on broad, high-level indexing terms such as Medical Subject Headings (MeSH) or author keywords. These generic terms lack the semantic and technological resolution needed to track specific interventional modalities, pentaspline catheter architectures, novel safety concerns such as microvascular spasm or hemolysis, or subtle shifts in risk stratification scoring algorithms across consecutive publication years^18^.

Addressing this methodological limitation is of paramount clinical and scientific importance. As interventional cardiology and cardiac electrophysiology adopt novel therapeutic platforms at an accelerated rate, understanding the pace and nature of evidence dissemination is essential for clinical practice, health policy, and trial design. Quantitative tracking of immediate post-publication research reception allows clinicians, guideline committees, and investigators to identify emerging therapeutic trends, recognize early post-market safety signals, and evaluate how rapidly novel interventional strategies displace established standard-of-care treatments ^19^. Establishing a standardized, unbiased framework capable of dissecting granular scientific concepts in contemporary literature is therefore vital for providing an accurate, real-time lens into the evolving landscape of cardiovascular medicine.

To resolve these challenges, this study developed an objective analytical framework that combines OpenAlex topic indexing^20^, a fixed 18-month post-publication citation observation window^17^, and multi-tier large language model (LLM)--assisted concept mining with canonical taxonomy normalization^21^. By evaluating the 50 most cited original research articles per publication year within OpenAlex topic T10065 across two consecutive cohorts—the Class of 2025 (papers published in 2023) and the Class of 2026 (papers published in 2024) —this study aimed to track temporal shifts in normalized concept prevalence and identify emerging thematic paradigms in high-impact atrial fibrillation management and outcomes research.

## METHODS

### Study Design and Data Source

This study is a retrospective, comparative bibliometric analysis of highly cited research on atrial fibrillation management and outcomes. The analytical framework integrates literature metadata from OpenAlex^22,23^ with large language model (LLM)--assisted curation and concept mining^24^ to characterize how influential papers and their underlying research themes differ between two publication cohorts. Papers published in calendar year 2023 constitute the Class of 2025 cohort, and papers published in calendar year 2024 constitute the Class of 2026 cohort, reflecting the elapsed time required for post-publication citation accrual before leaderboard assembly.

Bibliographic records were drawn from OpenAlex topic T10065 (Atrial Fibrillation Management and Outcomes), which indexes scholarly works tagged to this research area within the OpenAlex topic taxonomy. The overall workflow proceeded from (1) identification and ranking of the most cited on-topic research articles within each cohort, (2) extraction of paper-level scientific concepts from titles and abstracts, (3) normalization of extracted terms into a shared canonical taxonomy and higher-level parent themes spanning both cohorts, and (4) quantitative comparison and visualization of concept prevalence across cohorts.

### Identification of Highly Cited Papers

To identify the most influential contemporary research within the topic, papers were ranked by the number of citations received within a fixed 18-month post-publication window rather than by lifetime citation count^17^. For each candidate work, the 18-month citation count was defined as the number of distinct citing publications in OpenAlex whose publication dates fell on or after the target paper’s publication date and on or before the date 18 months later (inclusive). Only papers for which the full 18-month observation period had elapsed were eligible for window-based citation scoring. Candidate retrieval within each cohort was restricted to OpenAlex record type article and to works assigned to topic T10065 with publication dates spanning the corresponding calendar year (1 January to 31 December).

Exact top-50 retrieval by 18-month citations was performed using a threshold-expansion procedure designed to minimize expensive per-paper citation-window queries while preserving ranking correctness^25^. An initial candidate set was formed from the top papers by total lifetime citations, and 18-month citation counts were computed for eligible works in that set. The 18-month citation count of the 50th-ranked paper in this initial set defined a threshold; all papers in the topic with total lifetime citations at or above that threshold were then retrieved and scored for 18-month citations. The final leaderboard comprised the 50 highest-ranked papers by 18-month citations, with total lifetime citations used as a tie-breaker when 18-month counts were equal.

Because OpenAlex topic assignment and bibliographic metadata do not guarantee semantic homogeneity at the level of individual study questions, each paper in the oversized candidate pool was further evaluated for topical relevance and article type using an LLM classifier applied to the title and abstract^26,27^. A paper was retained only if the classifier judged its research to address atrial fibrillation management and outcomes and if it represented original research (including primary clinical trials and methods or software contributions), excluding reviews, meta-analyses, editorials, and commentaries. From the pool of on-topic research articles, the 50 papers with the highest 18-month citation counts in each cohort were selected for downstream concept analysis.

### Concept Extraction

Scientific concepts were extracted independently from each selected paper’s title and abstract using an LLM prompt designed for literature concept mining^28^. The extraction task requested concise concept labels representing paper-specific scientific content, including named entities, methods, technologies, populations, interventions, phenotypes, and defined sub-problems central to the study, while discouraging generic field labels and routine study-design vocabulary unless such terms were central to the contribution. Each paper was limited to a maximum of 20 extracted concepts. When an abstract was unavailable or very short, classification and extraction relied on the title.

### Taxonomy Construction and Concept Normalization

Extracted concepts from both cohorts were consolidated into a unified taxonomy to enable cross-cohort comparison. Normalization proceeded in two stages applied to the combined set of unique extracted strings.

In the first stage, synonymous or closely related extracted terms were grouped under canonical concept names. An embedding model was used to cluster semantically similar terms ^29,30^, and an LLM assigned canonical labels to synonym groups within similarity-batched inputs. Canonical names not merged into multi-term groups were retained as singleton entries. This stage yielded 719 canonical concepts from 1,061 deduplicated extracted strings across 100 papers (50 per cohort).

In the second stage, each canonical concept was assigned to exactly one parent theme representing a broader thematic category within atrial fibrillation management and outcomes research. Parent assignment used the same embedding-assisted batching strategy and an LLM mapping prompt^31^, producing 32 parent themes. Every extracted concept was mapped through the canonical layer to its parent theme, yielding a three-level structure (original extraction, canonical concept, parent theme) for each paper.

### Temporal Comparison of Concept Frequencies

Concept prevalence was compared between cohorts at both the canonical and parent-theme levels^32^. For each concept, a paper-level presence count was computed separately for the Class of 2025 and Class of 2026 corpora: a concept contributed one count to a cohort if it appeared at least once among that paper’s normalized concepts, regardless of how many synonymous extractions mapped to the same canonical label within the paper. Normalized frequency was defined as the paper-level presence count divided by the number of papers in the cohort (50 per year).

For each concept, the frequency difference was calculated as the Class of 2026 normalized frequency minus the Class of 2025 normalized frequency. Descriptive fold change was computed as the ratio of Class of 2026 to Class of 2025 normalized frequencies with a small offset (10^-12^) added to both numerator and denominator to stabilize ratios when frequencies approached zero^33^; log_2 fold change was derived from this fold change. These comparisons were descriptive; no inferential hypothesis tests, significance thresholds, or multiple-comparison corrections were applied to concept frequency differences.

### Data Visualization

Temporal shifts in concept prevalence were visualized using three complementary figure types for each taxonomy level (canonical topics and parent themes)^34^. Diverging horizontal bar charts displayed concepts with the largest positive and negative frequency differences between cohorts (10 concepts per direction, deduplicated when overlap occurred). Scatter plots placed Class of 2025 normalized frequency on the horizontal axis and Class of 2026 normalized frequency on the vertical axis for concepts included in the comparative figure set at each taxonomy level, with a diagonal reference line indicating equal prevalence; concepts with the largest absolute frequency differences within that set were annotated to highlight cohort-specific shifts (up to six canonical topics and up to five parent themes). Grouped horizontal bar charts showed normalized frequencies in both cohorts for the 15 concepts with the highest Class of 2026 normalized frequency.

Word clouds were generated separately for each cohort to summarize dominant themes among the most cited papers^35^. For each publication year, the 50 highest-ranked papers by 18-month citations were used. Word size was proportional to the number of papers in that cohort in which each canonical topic or parent theme appeared at least once. Word clouds were rendered for canonical topics and parent themes using a discrete color palette with one color per displayed term.

### Statistical Software and Computational Tools

All data processing, tabulation, and figure generation were performed in Python 3.12. Tabular operations used pandas 3.0.3^36^. Static figures were produced with matplotlib 3.11.0 at 300 dots per inch ^37^; word clouds used wordcloud 1.9.6^35^.

The manuscript was authored and rendered using the K2F document engine (https://k2f.dev/; https://github.com/suzheng/k2f). Bibliographic retrieval and citation-window counting used the OpenAlex REST API^22^. LLM-based classification, concept extraction, normalization, and parent-theme assignment were conducted through configured API endpoints using claude-sonnet-5:floor for topic relevance screening and concept extraction and deepseek/deepseek-v4-pro for canonical and parent-theme mapping; text embeddings for normalization batching used google/gemini-embedding-001.

## RESULTS

### Leaderboard Composition and Citation Accrual

Bibliometric assembly identified 50 on-topic research articles per publication cohort ranked by 18-month citation counts within OpenAlex topic T10065 (Atrial Fibrillation Management and Outcomes). For the Class of 2025 cohort (papers published in 2023), 18-month citation counts ranged from 29 to 318 across the top 50 papers. The highest-ranked paper reported a randomized comparison of pulsed field ablation versus conventional thermal ablation for paroxysmal atrial fibrillation^9^. Additional highly cited Class of 2025 publications included pivotal anticoagulation trials in subclinical atrial fibrillation and atrial high-rate episodes^13,38^ catheter ablation in end-stage heart failure with atrial fibrillation^39^, and early safety and effectiveness reports from pulsed field ablation registries and pivotal trials^40,41^.

For the Class of 2026 cohort (papers published in 2024), 18-month citation counts among the top 50 papers ranged from 29 to 278. The leading paper described post-approval safety outcomes for pulsed field ablation in more than 17,000 patients (MANIFEST-17K)^10^. Other top-ranked Class of 2026 publications included randomized trials of andexanet for factor Xa inhibitor--associated intracerebral hemorrhage ^42^, left atrial appendage closure after ablation ^43^, global burden estimates for atrial fibrillation and atrial flutter ^44^, and a phase 3 comparison of asundexian versus apixaban^15^. Across both cohorts, the most cited works were predominantly multicenter clinical trials, large registry analyses, and population-based epidemiological studies published in general medical and cardiovascular journals.

The complete ranked lists of the top 50 on-topic papers for each cohort are provided in Supplementary Tables S1^1,9,13,38-41,45-87^ and S2^10,12,15,42-44,88-131^.

### Dominant Scientific Themes in Contemporary Research

A ranked overview of the ten canonical topics with the largest Class of 2026 normalized frequencies emphasized pulsed field ablation, catheter ablation, pulmonary vein isolation, CHA2DS2-VASc score, thromboembolism, radiofrequency ablation, anticoagulation therapy, persistent atrial fibrillation, ischemic stroke, and arrhythmia recurrence (Figure **1)**. Word clouds summarizing parent themes and canonical topics among the 50 highest-ranked Class of 2026 papers by 18-month citation count highlighted catheter-based intervention, ablation energy technologies, thromboembolic risk assessment, and anticoagulation as visually dominant research areas (Figure 2, Figure 3). At the parent-theme level, Catheter Ablation: Procedures and Techniques, Ablation Technologies and Energy Sources, Risk Stratification and Predictive Models, and Anticoagulation and Stroke Prevention occupied the largest display areas (Figure 2). At the canonical-topic level, pulsed field ablation, pulmonary vein isolation, catheter ablation, CHA2DS2-VASc score, radiofrequency ablation, thromboembolism, anticoagulation therapy, and ischemic stroke were among the most prominently displayed terms (Figure 3).

**Figure 1.**
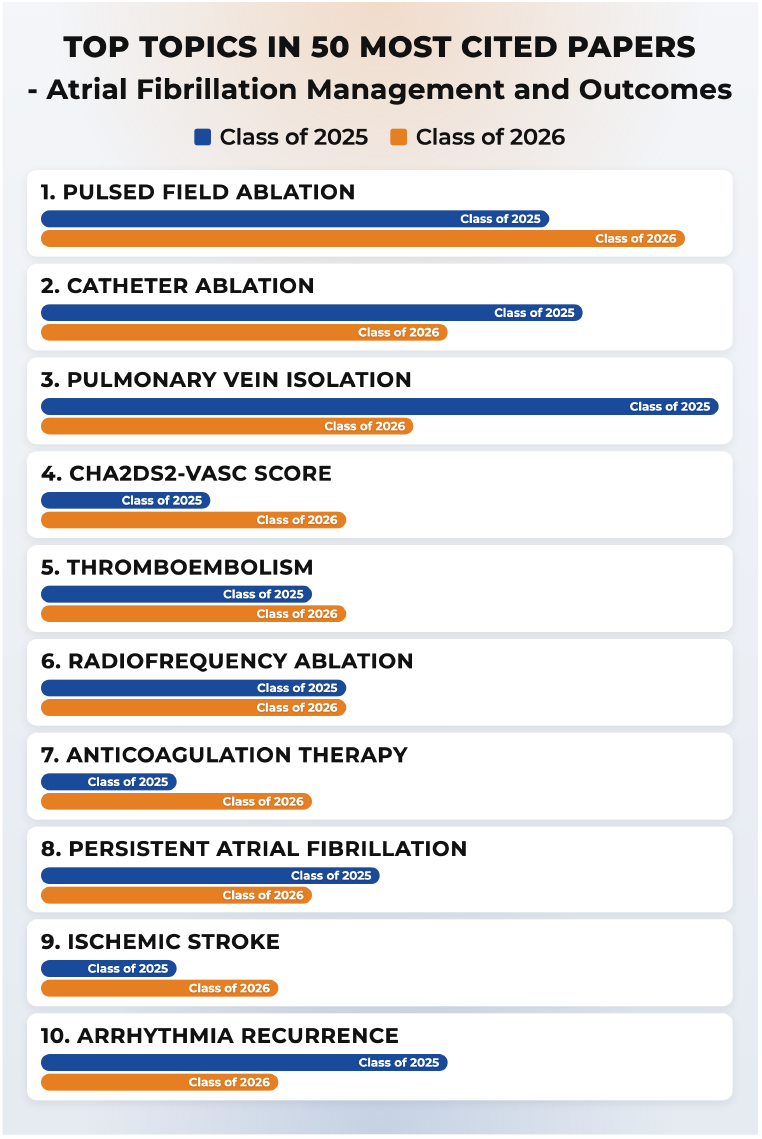
Top canonical topics in the 50 most cited Class of 2026 papers. Horizontal bar cards comparing normalized topic frequency between the Class of 2025 (blue bars) and Class of 2026 (orange bars) cohorts for the ten highest-ranked canonical topics in the Class of 2026 corpus. Topic labels appear as card headers; bar length is proportional to normalized frequency (paper-level presence count divided by cohort sample size n = 50). PFA, pulsed field ablation; PVI, pulmonary vein isolation; AF, atrial fibrillation.

**Figure 2.**
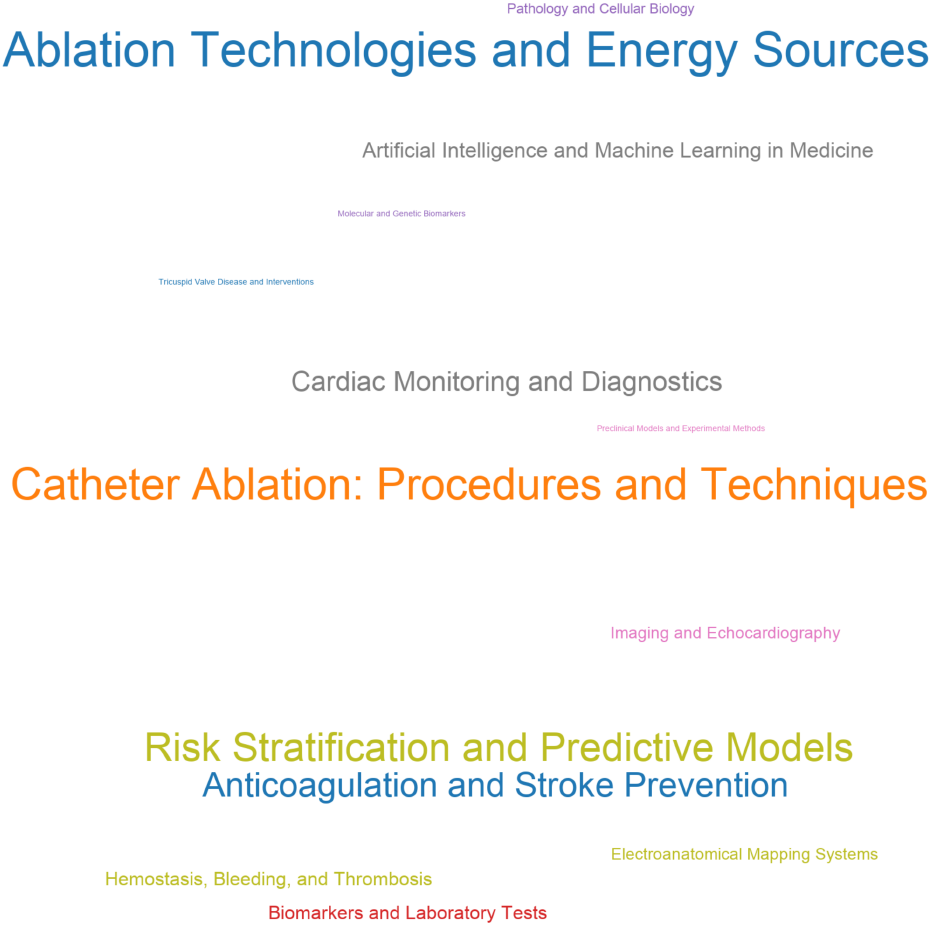
Parent-theme word cloud for the Class of 2026 cohort. Word cloud visualization of parent themes among the 50 highest-ranked Class of 2026 papers by 18-month citation count. Font size is proportional to the number of papers in which each parent theme appeared at least once across the cohort corpus. Terms are rendered in distinct colors without semantic cluster grouping.

**Figure 3.**
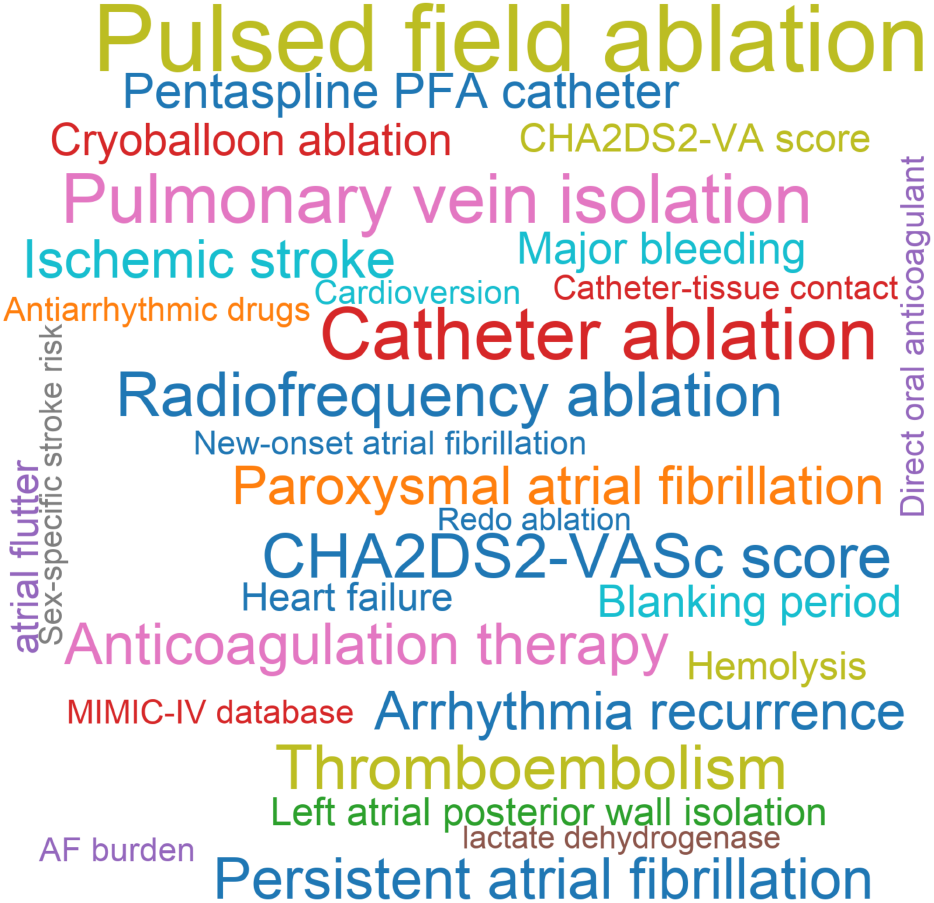
Canonical-topic word cloud for the Class of 2026 cohort. Word cloud visualization of granular canonical topics among the 50 highest-ranked Class of 2026 papers by 18-month citation count. Font size is proportional to paper-level presence frequency across the cohort corpus. Terms are rendered in distinct colors without semantic grouping. PFA, pulsed field ablation; PVI, pulmonary vein isolation; DOAC, direct oral anticoagulant.

### Temporal Shifts in Parent-Theme Prevalence

Comparison of normalized parent-theme frequencies between cohorts indicated redistribution of thematic emphasis while core ablation-related themes remained prevalent. Among parent themes represented in the comparative figure set, Risk Stratification and Predictive Models showed the largest positive frequency difference (normalized frequency 0.30 in the Class of 2025 cohort versus 0.38 in the Class of 2026 cohort; difference +0.08), followed by Ablation Technologies and Energy Sources (0.40 versus 0.46; +0.06), Hemostasis, Bleeding, and Thrombosis (0.06 versus 0.10; +0.04), and Anticoagulation and Stroke Prevention (0.26 versus 0.30; +0.04) (Figure 4). The largest negative difference was observed for Catheter Ablation: Procedures and Techniques (0.58 versus 0.46; -0.12), with additional relative declines for Electroanatomical Mapping Systems (0.14 versus 0.08; -0.06), Cardiac Monitoring and Diagnostics (0.24 versus 0.20; -0.04), and Imaging and Echocardiography (0.12 versus 0.08; -0.04) (Figure 4). Artificial Intelligence and Machine Learning in Medicine, Comorbidities and Associated Conditions, Preclinical Models and Experimental Methods, and Tricuspid Valve Disease and Interventions showed no change in normalized frequency between cohorts.

**Figure 4.**
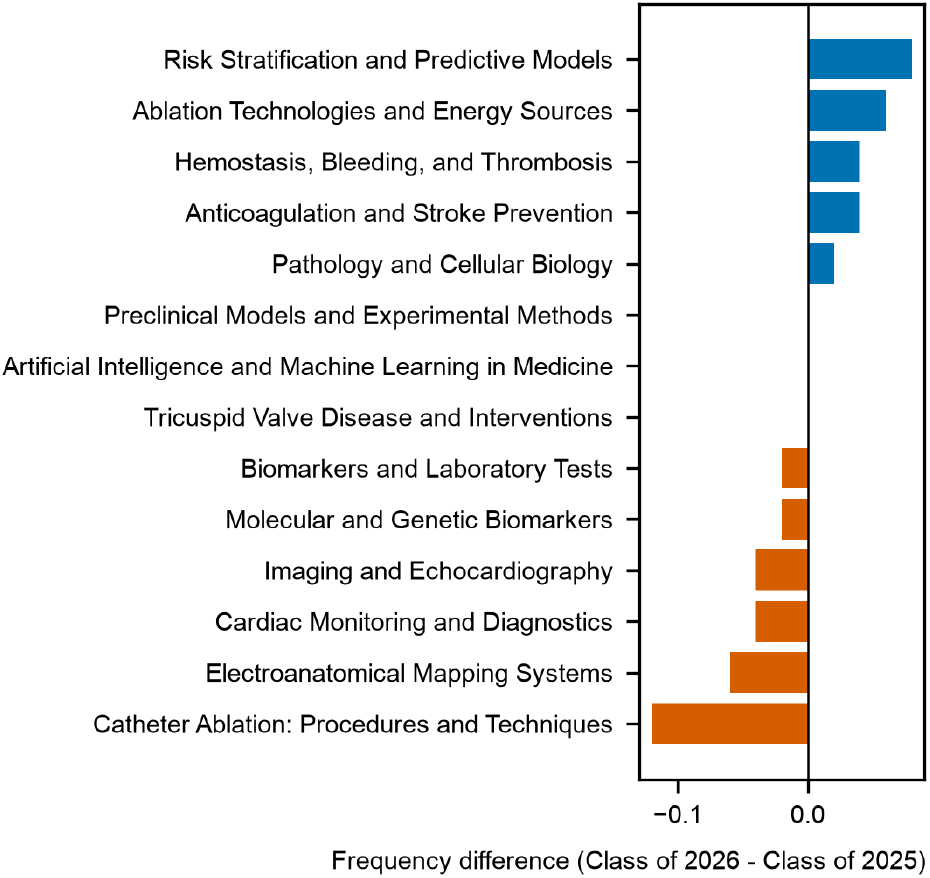
Parent-theme frequency difference between publication cohorts. Diverging horizontal bar chart displaying normalized frequency differences (Class of 2026 minus Class of 2025) for parent themes across the comparative figure set. Blue bars indicate positive frequency differences (increased prevalence in 2024); orange bars indicate negative frequency differences (decreased prevalence in 2024). Horizontal axis represents normalized frequency difference; vertical axis displays parent-theme labels.

Scatter analysis of parent-theme normalized frequencies placed most themes near the diagonal of equal cohort prevalence, while annotated outliers highlighted cohort-specific shifts (Figure 5). Catheter Ablation: Procedures and Techniques plotted below the equality line, consistent with lower r Class of 2026 prevalence. Risk Stratification and Predictive Models, Ablation Technologies and Energy Sources, and Hemostasis, Bleeding, and Thrombosis plotted above the equality line, indicating higher Class of 2026 prevalence. Electroanatomical Mapping Systems plotted below the equality line (Figure 5).

**Figure 5.**
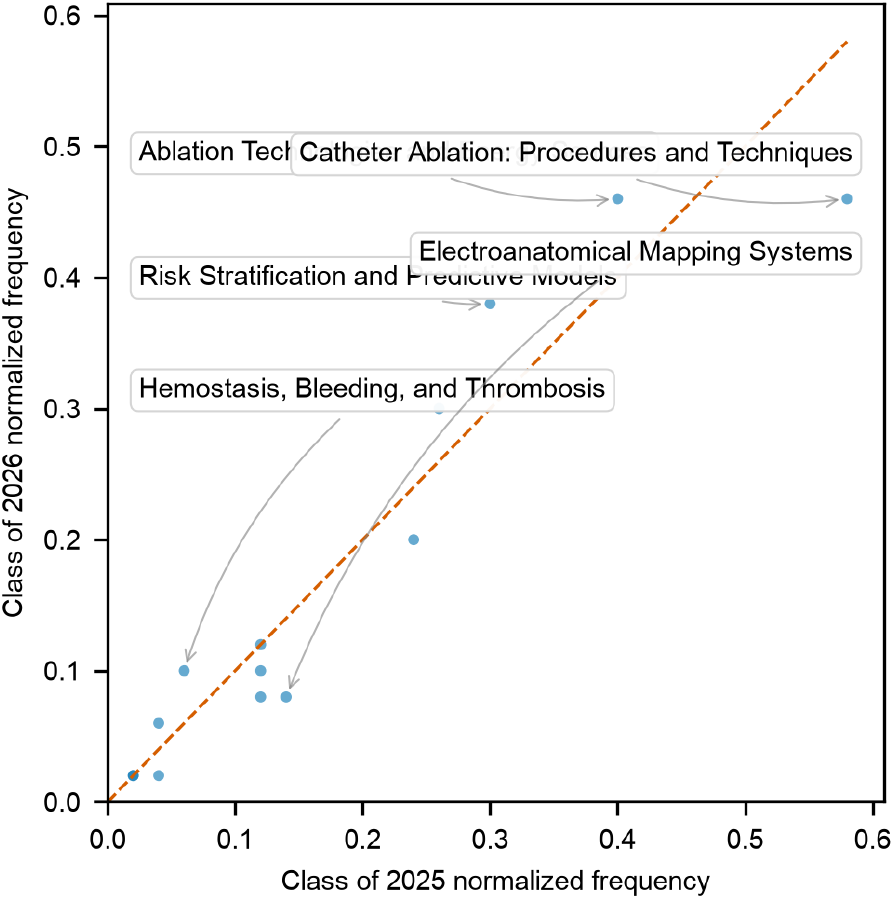
Parent-theme cohort scatter plot. Bivariate scatter plot comparing Class of 2025 (horizontal axis) versus Class of 2026 (vertical axis) normalized parent-theme frequencies. Each plotted circular point represents one parent theme. The dashed diagonal reference line indicates equal prevalence between cohorts. Annotated textual callouts highlight themes demonstrating the largest absolute frequency differences between publication years.

Grouped bar charts of the 14 parent themes with the highest Class of 2026 normalized frequencies showed that Catheter Ablation: Procedures and Techniques and Ablation Technologies and Energy Sources remained the two most frequent themes in both cohorts despite the relative decline in procedural emphasis (Figure 6). Risk Stratification and Predictive Models ranked third in the Class of 2026 cohort (normalized frequency 0.38), exceeding its Class of 2025 value (0.30). Anticoagulation and Stroke Prevention (0.30 versus 0.26) and Hemostasis, Bleeding, and Thrombosis (0.10 versus 0.06) also showed higher Class of 2026 frequencies, whereas Electroanatomical Mapping Systems (0.08 versus 0.14) and Cardiac Monitoring and Diagnostics (0.20 versus 0.24) showed lower Class of 2026 frequencies (Figure 6). Complete parent-theme counts and normalized frequencies for all 32 themes are reported in (Supplementary Table S3).

**Figure 6.**
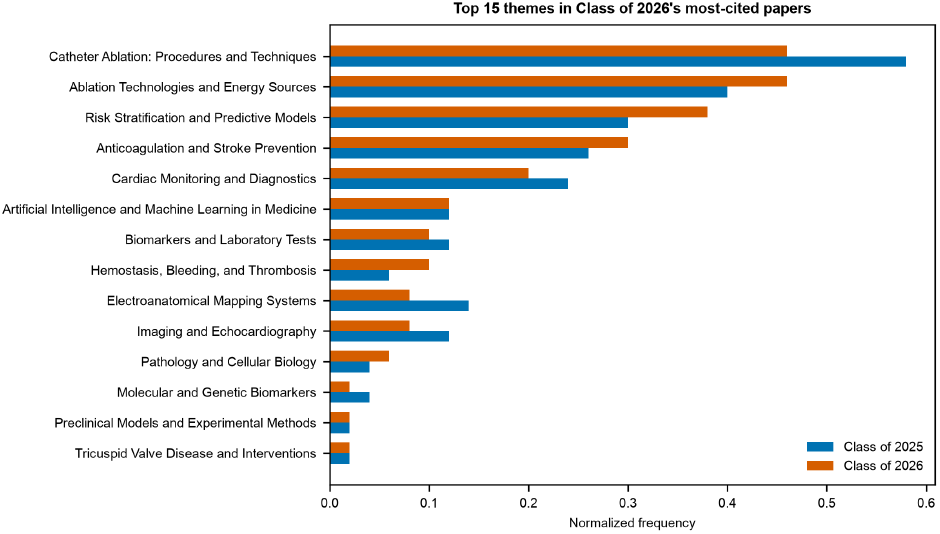
Top parent themes in Class of 2026 highly cited papers. Grouped horizontal bar chart of normalized frequencies for the 14 parent themes with the highest Class of 2026 prevalence. Blue bars denote the Class of 2025 cohort; orange bars denote the Class of 2026 cohort. Horizontal axis represents normalized frequency (paper presence count divided by cohort sample size n = 50).

### Temporal Shifts in Canonical-Topic Prevalence

At the canonical-topic level, pulsed field ablation showed the largest positive frequency difference among topics in the comparative figure set (normalized frequency 0.30 in the Class of 2025 cohort versus 0.38 in the Class of 2026 cohort; +0.08), alongside increases for CHA2DS2-VASc score (0.10 versus 0.18; +0.08), anticoagulation therapy (0.08 versus 0.16; +0.08), and CHA2DS2-VA score (0.00 versus 0.08; +0.08) (Figure 7). Topics that appeared only in the Class of 2026 cohort at normalized frequency 0.06 included new-onset atrial fibrillation, MIMIC-IV database, and lactate dehydrogenase (+0.06 each). The largest negative differences were observed for pulmonary vein isolation (0.40 versus 0.22; -0.18), paroxysmal atrial fibrillation (0.28 versus 0.14; -0.14), arrhythmia recurrence (0.24 versus 0.14; -0.10), catheter ablation (0.32 versus 0.24; -0.08), antiarrhythmic drugs (0.14 versus 0.06; -0.08), direct oral anticoagulant (0.14 versus 0.06; -0.08), cryoballoon ablation (0.18 versus 0.10; -0.08), major bleeding (0.18 versus 0.10; -0.08), and intracranial hemorrhage (0.12 versus 0.04; -0.08) (Figure 7). Additional Class of 2026 increases included ischemic stroke (0.08 versus 0.14; +0.06), hemolysis (0.02 versus 0.08; +0.06), and heart failure (0.02 versus 0.08; +0.06).

**Figure 7.**
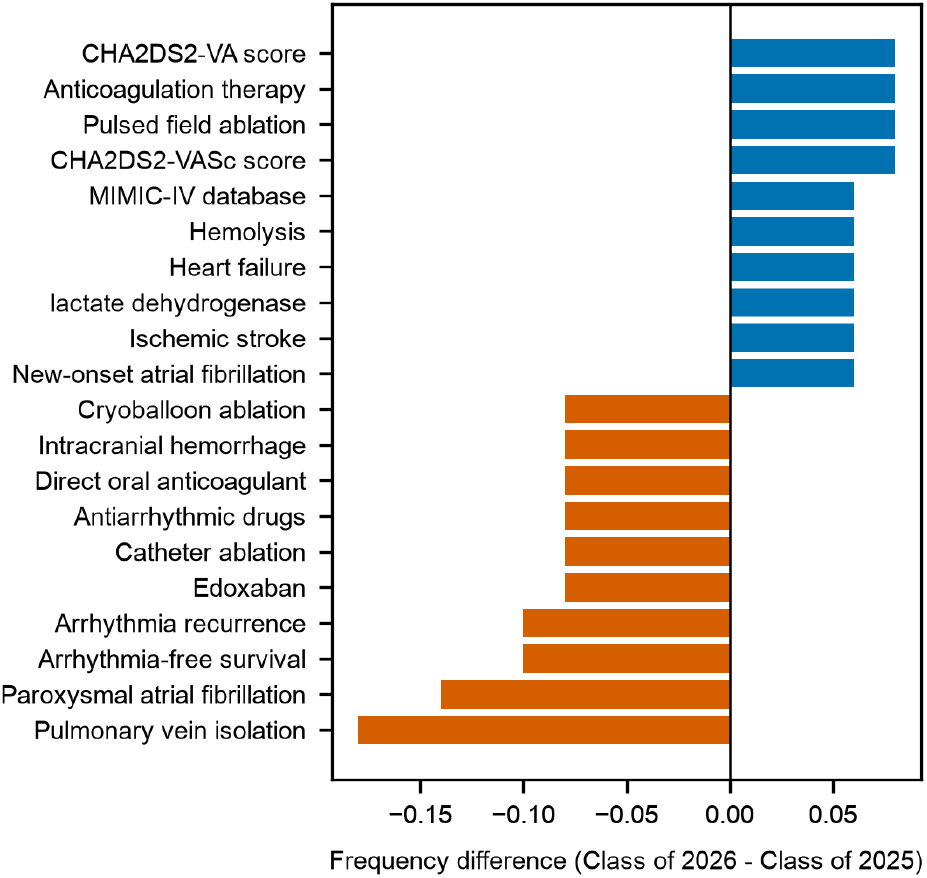
Canonical-topic frequency difference between publication cohorts. Diverging horizontal bar chart displaying normalized frequency differences (Class of 2026 minus Class of 2025) for canonical topics in the comparative figure set. Blue bars represent positive frequency differences; orange bars represent negative frequency differences. Horizontal axis indicates normalized frequency difference; vertical axis displays canonical-topic labels. PFA, pulsed field ablation; PVI, pulmonary vein isolation; DOAC, direct oral anticoagulant; AF, atrial fibrillation.

Scatter analysis of canonical topics showed pulsed field ablation, anticoagulation therapy, and CHA2DS2-VASc score above the equality line, whereas pulmonary vein isolation, paroxysmal atrial fibrillation, arrhythmia recurrence, and arrhythmia-free survival plotted below the equality line (Figure 8). Grouped bars for the 15 canonical topics with the highest Class of 2026 normalized frequencies confirmed pulsed field ablation as the leading topic (0.38), followed by catheter ablation (0.24), pulmonary vein isolation (0.22), CHA2DS2-VASc score (0.18), thromboembolism (0.18), radiofrequency ablation (0.18), persistent atrial fibrillation (0.16), anticoagulation therapy (0.16), paroxysmal atrial fibrillation (0.14), ischemic stroke (0.14), arrhythmia recurrence (0.14), pentaspline PFA catheter (0.12), major bleeding (0.10), blanking period (0.10), and cryoballoon ablation (0.10) (Figure 9). Complete canonical-topic counts and normalized frequencies for all 719 topics are reported in (Supplementary Table S4).

**Figure 8.**
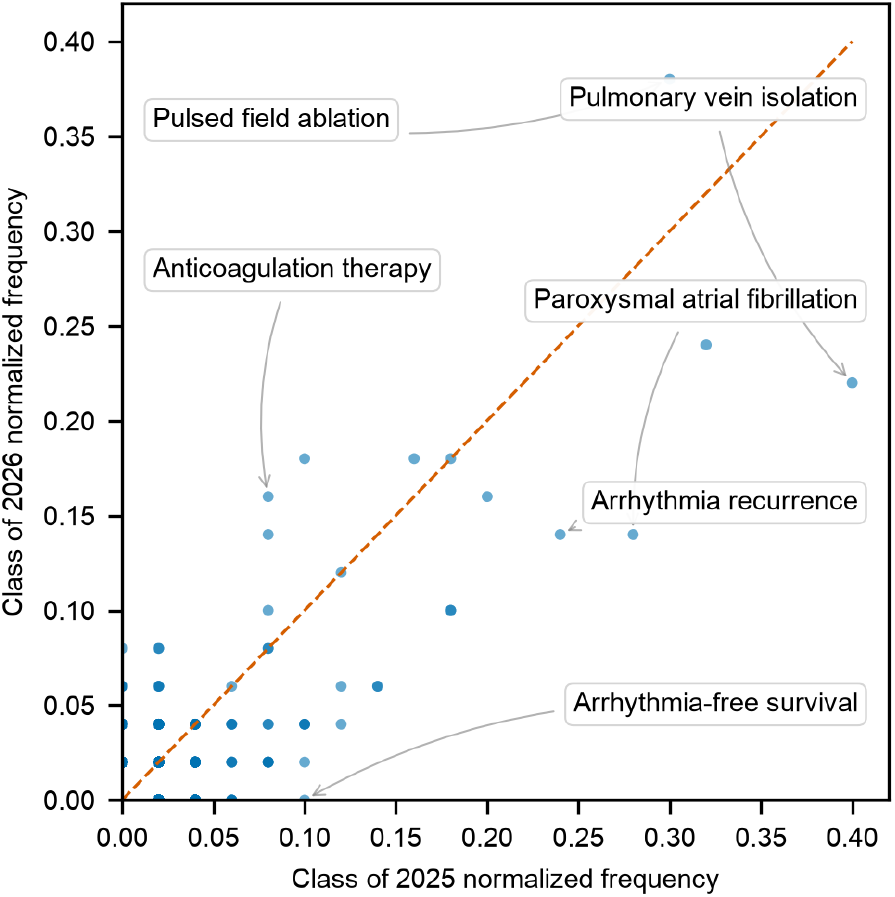
Canonical-topic cohort scatter plot. Bivariate scatter plot comparing Class of 2025 (horizontal axis) versus Class of 2026 (vertical axis) normalized canonical-topic frequencies. Each point represents an individual canonical topic. The dashed diagonal reference line indicates equal cohort frequency. Annotated labels highlight outlying topics exhibiting substantial frequency shifts between cohorts.

**Figure 9.**
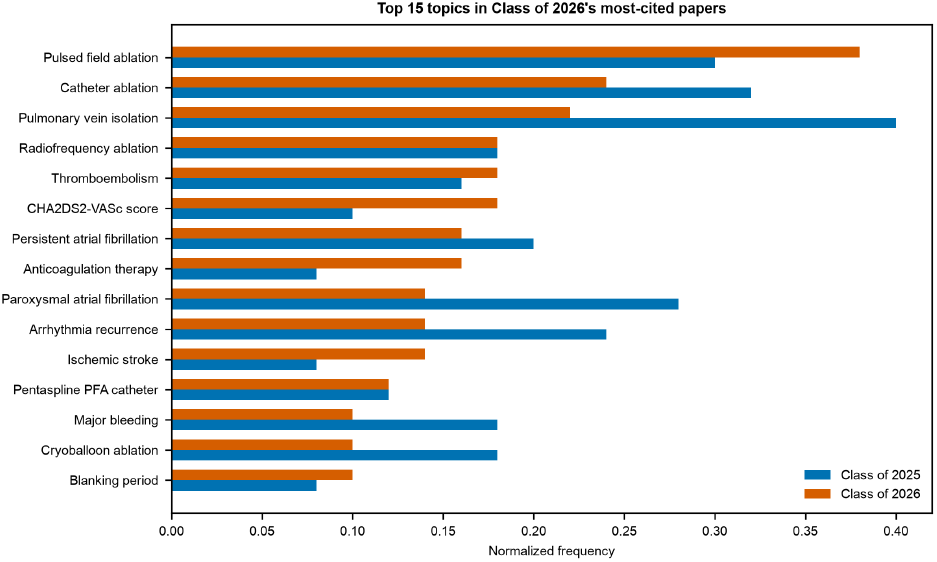
Top canonical topics in Class of 2026 highly cited papers. Grouped horizontal bar chart of normalized frequencies for the 15 canonical topics with the highest Class of 2026 prevalence. Blue bars represent the Class of 2025 cohort; orange bars represent the Class of 2026 cohort. Horizontal axis represents normalized frequency (paper presence count divided by cohort sample size n = 50). PFA, pulsed field ablation; PVI, pulmonary vein isolation; DOAC, direct oral anticoagulant.

## DISCUSSION

This comparative bibliometric and concept-mining analysis reveals a rapid shift in contemporary atrial fibrillation research between the Class of 2025 (2023 publications) and Class of 2026 (2024 publications) cohorts. Across the 100 highest-cited original research articles evaluated within OpenAlex topic T10065, electrophysiological ablation technologies and stroke prevention strategies maintained overarching dominance as parent research themes. However, granular concept tracking demonstrated a marked redistribution of scientific attention within these domains.

Pulsed field ablation emerged as the single most prevalent canonical topic in the 2024 cohort, displacing conventional thermal modalities such as radiofrequency ablation and cryoballoon ablation^9,40,41^ Concurrently, procedural focus shifted from basic pulmonary vein isolation mechanics toward technology-specific safety, lesion durability, and complication profiling, including intravascular hemolysis and microvascular spasm^98,132^. In stroke risk management, research emphasis shifted toward refining risk stratification algorithms—highlighted by comparisons between the CHA2DS2-VASc score and non-sex CHA2DS2-VA score^12,95^—as well as expanding indications for anticoagulation in subclinical atrial fibrillation ^13,38^ evaluating Factor Xa reversal strategies^42^, and exploring novel Factor XIa inhibitors^15,16.^

The prominent surge in pulsed field ablation prevalence reflects a major paradigm shift in interventional electrophysiology, where non-thermal tissue-selective electroporation is rapidly displacing traditional thermal modalities. Pivotal trial literature comparing pulsed field ablation with thermal ablation^9^ and single-arm registry reports^40,41^ demonstrated non-inferior procedural efficacy alongside superior procedural speed and reduced collateral tissue injury. Highly cited literature in the subsequent cohort moved beyond initial feasibility to evaluate real-world safety in over 17,000 patients^10^ and post-procedural complication management.

The corresponding rise of canonical topics such as hemolysis, coronary artery spasm, and acute kidney injury reflects a clinical maturation, as high-frequency energy delivery and pentaspline catheters have been linked to transient hemolysis and coronary vasoconstriction requiring targeted pharmacological mitigation^98,132^. This procedural evolution explains frequency declines for generalized pulmonary vein isolation and paroxysmal atrial fibrillation, as electrophysiology research matured toward addressing persistent atrial fibrillation substrates, lesion durability, and re-evaluating post-ablation blanking period definitions^40,108^.

In parallel, temporal redistribution within stroke prevention underscores growing clinical focus on personalized risk modeling and targeted antithrombotic therapies. The scientific focus on risk stratification was driven by critical re-evaluations of the CHA2DS2-VASc score versus the non-sex CHA2DS2-VA scheme^12,95^. This debate addresses whether female sex functions as an independent risk modifier or an age-dependent risk amplifier, with large population cohorts demonstrating that excluding sex-category scoring provides comparable or superior discrimination for ischemic stroke risk in modeen anticoagulant--treated populations ^127^.

Furthermore, expanding anticoagulation literature reflects trial results in subclinical atrial fibrillation and atrial high-rate episodes, where direct oral anticoagulant therapy reduced ischemic stroke at the expense of increased major bleeding^13,38^. Concurrently, clinical focus expanded to hemorrhage management—highlighted by randomized evidence supporting andexanet alfa for Factor Xa inhibitor--associated intracranial hemorrhage^42^—and next-generation Factor XIa inhibitors such as asundexian^15,16^ —to decouple antithrombotic efficacy from bleeding risk.

Comparing these results with historical bibliometric analyses of atrial fibrillation research highlights both persistent core priorities and unprecedented technological shifts. Earlier bibliometric mapping of cardiovascular electrophysiology from 1980 through 2021 demonstrated that catheter ablation and oral anticoagulation formed the twin pillars of atrial fibrillation literature, with radiofrequency energy and cryoballoon systems dominating procedural publications^133^. However, whereas earlier technological transitions—such as cryoballoon adoption over radiofrequency ablation—unfolded gradually across more than a decade^134^, current data reveal exceptionally swift citation displacement, with pulsed field ablation achieving primary thematic leadership within 18 to 24 months of pivotal trial publication^9,10.^

Furthermore, while previous literature focused heavily on long-term recurrence rates following thermal pulmonary vein isolation^135^, top-cited attention rapidly pivoted to pulsed field ablation--specific biological interactions, such as microvascular spasm and red blood cell lysis^98,132^. This contrast underscores how modern electrophysiology assimilates innovative interventional modalities at a substantially faster pace than in prior decades.

The novelty of this study lies in its objective framework combining a fixed 18-month citation accrual window with multi-tiered, large language model--assisted concept mining to capture high-impact medical paradigm shifts in real-time. Traditional bibliometric evaluations rely on lifetime citation counts, which inherently bias results toward older publications and obscure rapid research evolution^136,137^. By enforcing a strict 18-month post-publication observation window for consecutive calendar cohorts, this study isolates immediate post-publication scientific reception free from time-in-field bias.

Additionally, while standard bibliometric tools depend on author keywords or generic Medical Subject Headings that lack granular technological detail, our two-stage normalization pipeline extracted 1,061 granular concept strings and structured them into 719 canonical topics and 32 parent themes1^38^. This approach allowed for automated tracking of subtle semantic shifts—such as the transition from generic pulmonary vein isolation terms to specific catheter architectures like pentaspline pulsed field ablation systems—offering a novel computational methodology for literature synthesis.

Scientific and translational implications extend directly into clinical practice guidelines, electrophysiology laboratory workflows, and trial design. Rapid integration of pulsed field ablation into top-cited literature signals an urgent need for updated clinical guidelines detailing energy-specific procedural protocols, patient selection criteria, and post-ablation surveillance^12,139^. Translating pulsed field ablation into routine clinical care requires dedicated monitoring protocols for non-thermal complications, such as screening for intravascular hemolysis and periprocedural administration of coronary vasodilators during complex ablation^98,132^.

In stroke prevention, shifting emphasis toward non-sex CHA2DS2-VA scoring and subclinical atrial fibrillation management suggests that future clinical decision tools must incorporate nuanced, threshold-based risk algorithms rather than binary stroke risk scores^12,13,38^. Moreover, the rise of left atrial appendage closure trials following ablation^43^ points toward integrated pathways combining non-thermal ablation with mechanical stroke prophylaxis to minimize lifelong anticoagulation burden.

This study possesses key strengths as well as inherent limitations that warrant balanced consideration. A major strength is the rigorous methodology, combining automated bibliographic retrieval from OpenAlex^138^ with large language model filtering to exclude non-original research articles, ensuring concept frequencies reflected primary clinical evidence rather than review commentary. Two-tier embedding-assisted clustering and canonical taxonomy mapping provided a reproducible structure across publication years.

However, several limitations must be acknowledged. First, restricting analysis to the top 50 cited papers per cohort selectively highlights high-impact clinical trials and registry studies published in premier journals, potentially under-representing niche basic science or early-phase engineering innovations. Second, although the 18-month citation window standardizes evaluation time, citation counts remain a surrogate measure of academic interest rather than direct clinical utility^137^.

Third, concept extraction relied on titles and abstracts, which may omit detailed methodological nuances present only in full manuscript texts. Finally, comparative frequency shifts between cohorts were analyzed descriptively without inferential statistical testing.

Several critical research directions arise naturally from these observations. First, as pulsed field ablation expands globally, long-term post-market registries and pragmatic randomized trials are needed to establish whether non-thermal ablation translates into superior multi-year freedom from atrial arrhythmias compared with high-power short-duration radiofrequency ablation^140^. Second, basic and translational studies must further elucidate physiological mechanisms underlying pulsed field ablation--induced hemolysis, microvascular spasm, and endothelial response across diverse tissue geometries and pulse waveforms^98,132^.

Third, in pharmacological stroke prevention, long-term clinical outcomes from ongoing phase 3 trials evaluating Factor XI/XIa inhibitors must be monitored to determine if novel agents overcome the bleeding liabilities of Factor Xa inhibitors highlighted in recent trial literature^15,16^ Fourth, future bibliometric research should extend this concept-mining framework across multi-year longitudinal windows and integrate full-text parsing to trace how artificial intelligence algorithms and wearable device monitoring translate into real-world stroke risk reduction^141,142^.

In conclusion, this comparative bibliometric analysis demonstrates a dynamic transformation in highly cited atrial fibrillation research, characterized by rapid dominance of pulsed field ablation over conventional thermal modalities and a renewed focus on precision stroke risk stratification^9^,^12.^ By tracking normalized concept frequencies across consecutive publication cohorts within a fixed citation window, this study highlights how contemporary electrophysiology is pivoting from basic procedural feasibility toward non-thermal energy safety profiling, targeted complication management, and personalized antithrombotic strategies ^10,13^. As non-thermal ablation platforms and novel anticoagulation pathways continue to evolve, continuous quantitative mapping of scientific literature provides an essential lens for clinicians and researchers seeking to navigate emerging paradigms in cardiovascular medicine.

## CONCLUSIONS

This study demonstrates a rapid, measurable transformation in contemporary high-impact atrial fibrillation research between consecutive publication cohorts. Fixed-window citation tracking combined with LLM-assisted concept mining captured a clear technological pivot toward pulsed field ablation as the predominant catheter ablation modality, displacing conventional thermal pulmonary vein isolation. Concurrently, stroke prevention paradigms evolved from binary anticoagulation toward refined predictive scoring and non-thermal procedural safety profiling. These findings highlight both the dynamic trajectory of clinical electrophysiology and the power of fixed-window canonical concept mining for tracking real-time scientific evolution.

## Acknowledgments

The authors thank the OpenAlex team for providing open-access scholarly metadata and bibliometric resources.

## Data Availability

Source code and algorithm documentation are available at https://github.com/pepkio/pri-top-papers. Online explorer of the result is available at https://pri.pepkio.com

## Supplementary Information

**Supplementary Table S1**. Complete leaderboard of the 50 highest-cited Class of 2025 (2023 publication cohort) papers in OpenAlex topic T10065 ranked by 18-month post-publication citations, with OpenAlex IDs, publication metadata, total lifetime citations, and 18-month citation counts.

**Supplementary Table S2**. Complete leaderboard of the 50 highest-cited Class of 2026 (2024 publication cohort) papers in OpenAlex topic T10065 ranked by 18-month post-publication citations, with OpenAlex IDs, publication metadata, total lifetime citations, and 18-month citation counts.

**Supplementary Table S3**. All 32 parent themes with paper-level presence counts in each cohort, normalized frequencies, frequency difference (Class of 2026 minus Class of 2025), fold change, and log_2 fold change.

**Supplementary Table S4**. All 719 canonical topics with paper-level presence counts in each cohort, normalized frequencies, frequency difference (Class of 2026 minus Class of 2025), fold change, and log_2 fold change.

